# SpiDec: Computing Binodals and Interfacial Tension of Biomolecular Condensates From Simulations of Spinodal Decomposition

**DOI:** 10.1101/2022.06.15.496322

**Authors:** Konstantinos Mazarakos, Ramesh Prasad, Huan-Xiang Zhou

## Abstract

Phase separation of intrinsically disordered proteins (IDPs) is a phenomenon associated with many essential cellular processes, but a robust method to compute the binodal from molecular dynamics simulations of IDPs modeled at the all-atom level in explicit solvent is still elusive, due to the difficulty in preparing a suitable initial dense configuration and in achieving phase equilibration. Here we present SpiDec as such a method, based on spontaneous phase separation via spinodal decomposition that produces a dense slab when the system is initiated at a homogeneous, low density. After illustrating the method on four model systems, we apply SpiDec to a tetrapeptide modeled at the all-atom level and solvated in TIP3P water. The concentrations in the dense and dilute phases agree qualitatively with experimental results and point to binodals as a sensitive property for force-field parameterization. SpiDec may prove useful for the accurate determination of the phase equilibrium of IDPs.

**TOC Graphic:** 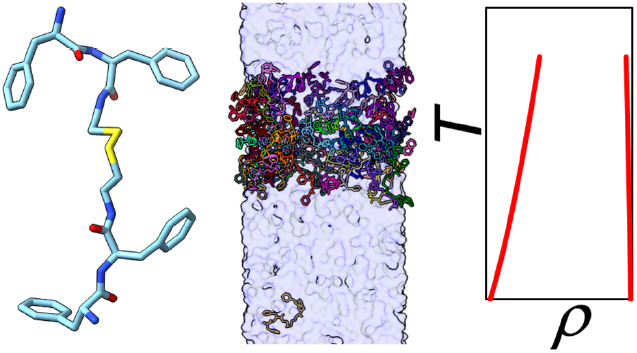

## 1. INTRODUCTION

Biomolecular condensates formed via liquid-liquid phase separation drive much of biology, but accurate computation of the binodal, representing the equilibrium concentrations in the bulk and dense phases as a function of temperature, based on atomistic modeling of the components presents a significant challenge. A related issue is the mechanism leading to phase separation. Theories predict that, depending on the initial densities or compositions, phase separation occurs by two mechanisms (Fig. 1). Inside the spinodal, the system is thermodynamically unstable and phase separation occurs spontaneously, in a process known as spinodal decomposition. This process is initiated by large-scale density fluctuations, leading to interconnected domains.

**Figure 1.**
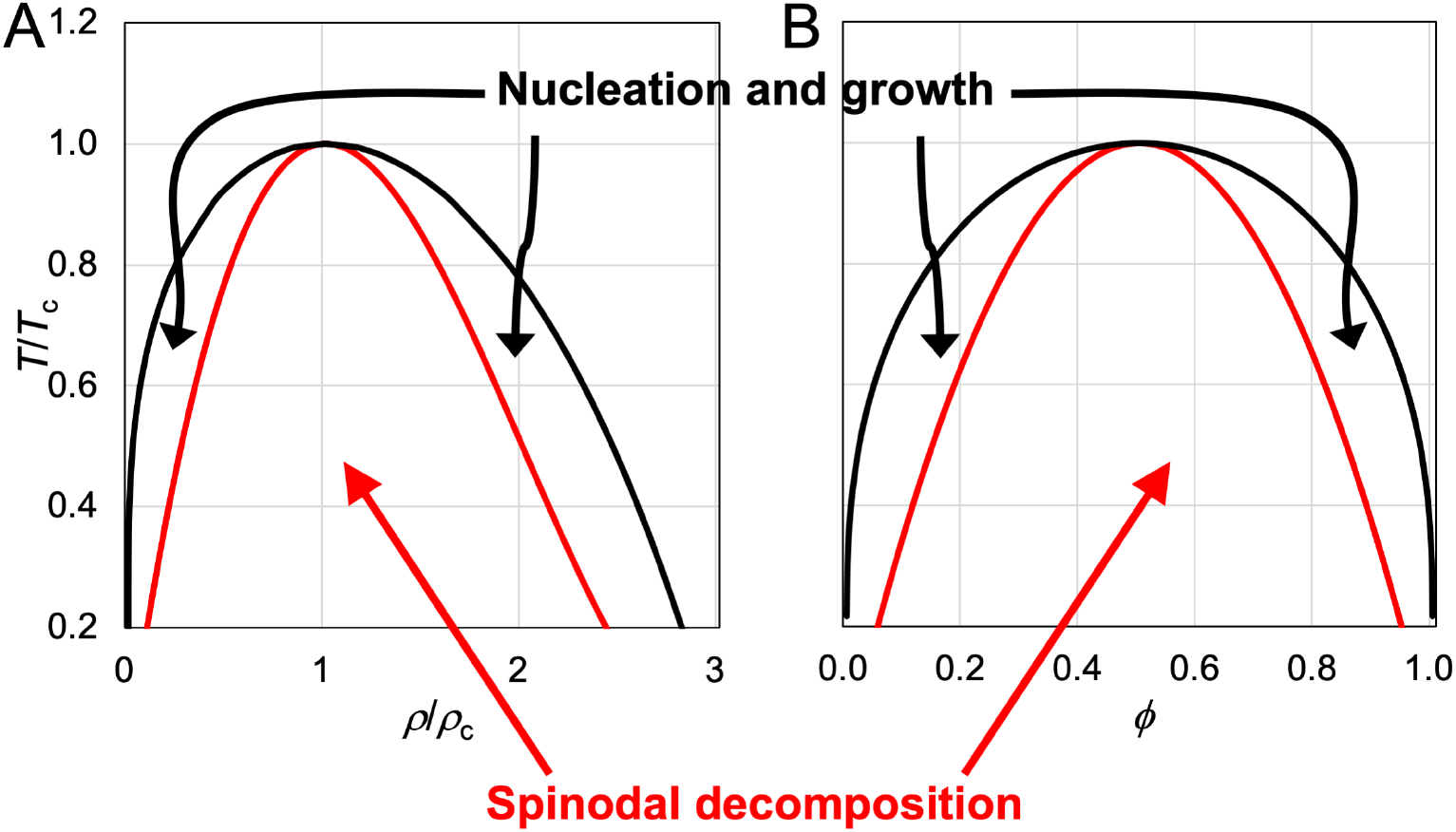
Binodals and spinodals of two model systems. (A) Van der Waals fluid, which satisfies the following equation of state: 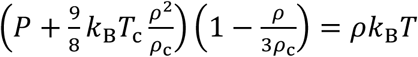 where *P, ρ, T* denote the pressure, density, and temperature, *T_c_* and *p_c_* denote the critical temperature and critical density, and *k_B_* is the Boltzmann constant. (B) Symmetric polymer blend that follows the Flory-Huggins theory for the Helmholtz free energy: 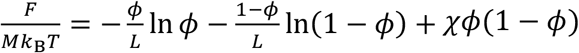, where *M* is the total number of polymer chains, *L* is the number of beads per chain, *ϕ* is the mole fraction of one polymer species in the binary blend, and *χ* measures the energy gap between inter- and intra-species interactions. Inside the spinodal region, the system is unstable and phase separates by spinodal decomposition; between the binodal and spinodal, the system is metastable and phase separates by nucleation and growth. Note that there has been some controversy regarding the location of the spinodal in computer simulations.^5–6^

Further condensation then leads to separated condensates. Between the binodal and spinodal, the system is metastable, and phase separation occurs by nucleation and growth. This mechanism is initiated by local density fluctuations, leading to the generation of nuclei. Further growth then produces stable condensates. The dividing line between these two mechanisms, i.e., the spinodal, is known for a number of theoretical models (Fig. 1) and has been measured for both structured and disordered proteins.^1–2^ Both spinodal decomposition and nucleation have been observed for the formation of biomolecular condensates.^2–3^ It has been suggested that cells may want to keep component concentrations to a minimum required for phase separation, i.e., crossing the low-concentration branch of the binodal just enough into the left metastable region, and thus nucleation and growth is the favored mechanism.^4^ However, in most theoretical models (Fig. 1), the spinodal covers the bulk of the area under the binodal, and therefore random initial preparations produce much higher chance for phase separation by spinodal decomposition than by nucleation and growth. Indeed, phase separation is observed instantaneously in many *in vitro* preparations; the rapid speed is consistent with spinodal decomposition.

In computer simulations of model systems, depending on the initial density, spinodal decomposition produces a variety of morphologies for the dense phase, including sphere, cylinder, slab, hollow cylinder, and hollow sphere.^5–6^ The slab morphology is of special interest because it represents the macroscopic form of the dense phase. More importantly, for this paper, the slab morphology is the basis of the method for computing binodals to be presented below.

Over the years, a variety of methods have been developed to compute the binodals of coarse-grained model systems (for a brief review and illustration, see ref. 7). One of the earliest methods is based on preparing a dense slab at the center of an otherwise empty simulation box.^8^ Molecular dynamics (MD) simulations then allow the dense and bulk phases to reach equilibration, and the densities of the two phases are then calculated to build the binodal. This classical slab method has now been applied to a variety of coarse-grained models of biomolecular condensates, from spherical particles and homopolymers (as crude models of structured proteins and IDPs)^9–10^ to residue-level models of IDPs^11–13^ to multi-bead-per-residue models of dipeptides.^14^ An exciting recent development is the migration of this method to simulations of all-atom models of IDPs in explicit solvent, by first carrying out full simulations at the coarse-grained level and then mapping to atomistic systems.^15–16^ Still, the mapping is not a trivial task and, after mapping, simulations at the all-atom level may fail to allow exchange of protein molecules between the phases, let alone reaching phase equilibrium.^15^ In comparison, Gibbs ensemble simulations,^17^ while proven useful for providing insights on biomolecular condensates^18–19^ and complex coacervates,^20–21^ are more restrictive, in particular due to the difficulty in inserting a polymer chain into a dense solution. Recent implementation by field-theoretic simulations has widened the applicability of the Gibbs ensemble,^22^ but application to all-atom models seems out of reach at present. Lastly the FMAP method, based on using the fast Fourier transform to evaluate intermolecular interaction energies, has been developed to calculate the binodals of structured proteins, but not IDPs, modeled at the all-atom level in implicit solvent.^23^

Here we present a method that we dub SpiDec, as an efficient alternative to the classical slab method. Instead of an initial dense slab, we start the system at a homogeneous, low density inside the spinodal. Spinodal decomposition then quickly brings the system to a slab morphology. We demonstrate this method both on four model systems and on a phase-separating tetrapeptide^24^ at the all-atom level in explicit solvent.

## 2. COMPUTATIONAL METHODS

### 2.1 Model Systems

We tested the SpiDec method on model systems composed of Lennard-Jones (LJ) particles, LJ chains, hydrophobic-hydrophilic (HP) chains, and patchy particles. The interaction potentials for LJ particles and LJ chains are the same as in our previous studies.^9, 25^ Specifically, the LJ particles interact via the LJ potential,

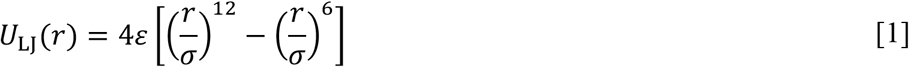

but with a cutoff imposed at *r_c_* = 3*σ*, so the actual potential function is shifted to

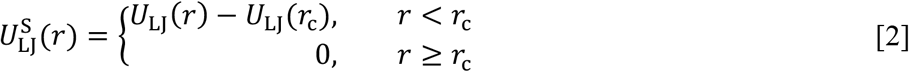

The LJ chains consist of 10 beads. The beads interact via a force-shifted LJ potential:

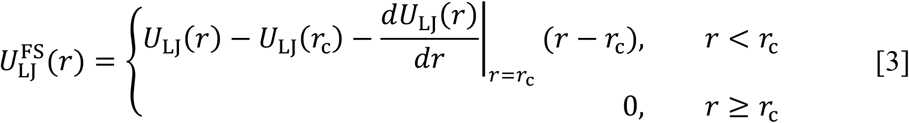

with a cutoff *r_c_* = 6*σ*. This potential is not applied to adjacent beads in a chain; instead they are connected by harmonic bonds with an equilibrium length of *σ* and a spring constant of 75,000*ε*/*σ*^2^. The HP chains are very much like the LJ chains, except that there are two kinds of beads, H and P.^13^ Each chain has two P beads, at positions 1 and 5; the rest are H beads. H beads interact with each other via the above force-shifted LJ potential, whereas all other pairwise interactions (H-P and P-P) were the purely repulsive Weeks-Chandler-Anderson (WCA) potential,^26^

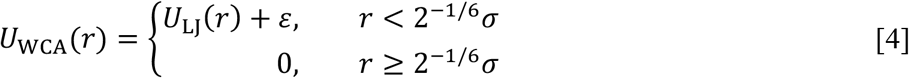

The interaction potential for patchy particles was the same as in our previous studies.^18–19^Each particle has four equal-sized circular patches with centers located at the vertices of a tetrahedron. Each patch has a spanning polar angle of *θ*_s_, which is set to a value (cos *θ*_s_ = 0.35) so the patches cover a fraction of 0.7 of the particle surface. The particles have a diameter *σ* and interact via the potential^27^

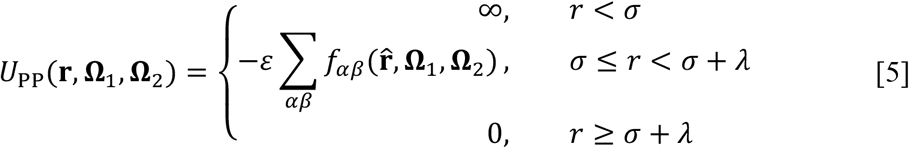

where the range of interaction, *λ*, is fixed to 0.5*σ*; 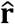 denotes the unit vector along the interparticle displacement **r** (pointing from particle 1 to particle 2); **Ω_*i*_** denotes the orientation of particle *i*; and 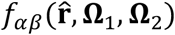 is 1 if patch *α* of particle 1 is “bonded” with patch *β* of particle 2, and 0 otherwise. The bonding condition is satisfied if the inter-particle vector falls within both patches, i.e., 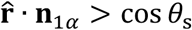 and 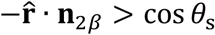, where n_*iα*_=is the unit vector from the center of particle *i* to the center of patch *α* on particle *i.* The orientation of each patchy particle was specified by the unit vectors along the three axes of a Cartesian coordinate system fixed to the particle.

### 2.2 Initialization for Simulations

We studied LJ particles by both MD and Monte-Carlo (MC) simulations, LJ and HP chains by MD simulations, and patchy particles by MC simulation. The simulation boxes were rectangular, with side length *L_x_* in two directions and *L_z_* (≥ *L_x_*) in the third direction. To prepare for MD simulations, particles or chains were randomly inserted into the simulation box at density *ρ_0_*. For MC simulations, the particles were initially placed on a cubic lattice spanning the simulation box, again at an initial density *ρ*_0_. The values of *ρ*_0_ are given below. The periodic boundary condition was applied during the simulations.

### 2.3 Ranges in Initial Density for Various Dense-Phase Morphologies

We scanned the initial density to identify the dense-phase morphologies of a given system, with the temperature at the lowest value (0.65, 1.70, 1.05, and 0.61 for LJ particles, LJ chains, HP chains, and patchy particles, respectively). The initial density was scanned up to 0.8, in two series. In the first series, the particle numbers were fixed (1000 for LJ particles and LJ and HP chains; 750 for patchy particles); the simulation boxes were cubic, and the side lengths were varied to span the range of initial densities, which was from 0.05 to 0.8 at increments of 0.05.

In the second series, done for LJ particles and LJ chains, *L_x_* was fixed (10 for LJ particles and 13 for LJ chains), and *L_z_/L_x_* was varied from 1 to 5 in increments of 0.25 or 0.5. The initial densities were 0.01 to 0.12 in increments of 0.01, 0.125 to 0.3 in increments of 0.025, 0.3 to 0.8 (for LJ particles) or 0.7 (for LJ chains) in increments of 0.05. The corresponding particle (or bead) numbers ranged from 10 to 4000 for LJ particles and from 20 to 7690 for LJ chains. For a given *L_z_/L_x_,* at increasing initial densities, spinodal decomposition or lack thereof results in seven distinct cases: a low-density homogenous phase, five dense-phase morphologies, and a high-density homogenous phase. For the boundary between the case of a low-density homogenous phase and the case of a dense phase with a spherical morphology, we took the midpoint between the highest initial density that resulted in a low-density homogenous phase and the lowest initial density that resulted in a dense phase with a spherical morphology. Sometimes it was uncertain whether to identify the status resulted from an intermediate initial density as a low-density homogenous phase or a dense phase with a spherical morphology; we then assigned that initial density as the boundary value. A similar procedure was followed to identify the next boundary, i.e., between a dense phase with a spherical morphology and a dense phase with a cylindrical morphology. The process continued until the last boundary, i.e., between a dense phase with a hollow sphere and a high-density homogenous phase, was determined.

### 2.4 Molecular Dynamics Simulations for LJ Particles and LJ and HP Chains

These simulations were carried out using the HOOMD-blue package (version 2.5.0) on GPUs.^28^ All the particles or beads have the same diameter *σ*, which sets the unit of lengths, and the same mass *m*. The units for number density, temperature, interfacial tension, and time are *σ*^-3^, *ε/k_B_, ε/σ*^2^, and 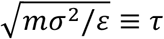, respectively. The MD simulations were carried out at constant particle (or bead) number *(N),* volume, and temperature. Temperature was regulated by the Langevin thermostat with a friction coefficient of *0.1m/τ.* The time step was 0.005τ for particle systems and 0.001τ for chain systems. For identifying the dense-phase morphologies of a given system, the initial densities were scanned over a range as described above. The presentation below applies to simulations where a slab morphology was formed and used to calculate binodals and interfacial tension.

The initial densities were 0.3 for LJ particles and 0.25 for LJ and HP chains. Separate simulations were carried out over a range of temperatures. To investigate the effects of system size and *L_z_/L_x_* ratio, we chose *N* in the range of 1000 to 10000, and *L_z_/L_x_* from 1.25 to 33.3 for LJ particles, from 1.16 to 18.2 for LJ chains, and 1.5 to 5 for HP chains. For LJ particles, the simulation length was 10 million steps, but was extended to 100 or 200 million steps when multiple slabs took a long time to fuse into a single slab (at *N* ≥ 6000 and *L_z_/L_x_* ≥ 20). The same was true for LJ chains, except that the longer simulations were 100 to 300 million steps and applied to more cases (*N* as low as 4000 and *L_z_/L_x_* as low as 2). The simulation length was 100 million steps for HP chains. The time interval for saving snapshots was 1000 time steps for simulations with a total length of 10 million steps and 10000 time steps for longer simulations.

### 2.5 Monte Carlo Simulations

For patchy particles, the initial density was 0.36, *N* ranged from 250 to 1250, and *L_z_/L_x_* was 1.5, 3, and 5. MC simulations were run for 2 million steps, and up to 5 million steps for larger *N* and elongated simulation boxes. Snapshots were saved for analysis once every 2000 MC steps. Each MC step consisted of either a displacement or a rotation (with equal probability) for every particle. The displacement was randomly selected inside a cube centered at the original position and with a side length of 0.09σ. The rotation was realized by picking an arbitrary new direction for the *z* axis of the particle-fixed coordinate system and rotating the line of nodes, defined as the cross product of the old and new *z* axes, by an angle to become the new *x* axis of the particle-fixed coordinate system.^19^ The latter was arbitrarily chosen between −0.05 to 0.05 radians. An MC simulation was also carried out for LJ particles at *ρ_0_* = 0.3, *N* = 1000, and *L_z_/L_x_* = 3 for 3 million steps (saving every 100 MC steps) to determine the binodal and interfacial tension, for comparison with the results obtained by MD simulations. In this case no rotation was included in the MC step.

### 2.6 Time for Phase Separation via Spinodal Decomposition

In simulations where the dense phase had a slab morphology, we monitored the time that it took for slabs to emerge from spinodal decomposition. In each saved snapshot, the density profile along the normal direction of the slabs (typically the *z* direction) was calculated in slices with a default thickness of 1σ and the maximum density was collected as a function of simulation time. After slabs were formed, the maximum densities reached a plateau. We took the first time that the maximum density exceeded the plateau value as the time to phase separate, denoted by τ_PS_. In some cases the default thickness of the slices for density calculation was adjusted so the resulting τ_PS_ was close to the value obtained by visual inspection, which looked for slabs with relatively flat surfaces.

When multiple slabs were formed, we also monitored the time, τ_ss_, that it took for the slabs to fuse one by one, finally into a single slab. The number of slabs in each saved snapshot after τ_PS_ was determined by the following procedure. Again, the density profile was calculated in slices of thickness 1σ. Each slice was then labeled as “H” (for high density) if its density exceeded a high cutoff *ρ*_H_, as “L” (for low density) if its density dipped below a low cutoff *ρ*_L_, or filtered out if its density fell between *ρ*_H_ and *ρ*_L_. In this way the density profile was converted to a sequence like HHHHLLL…HHH. Each transition from H to L or L to H in this sequence defined an interface. The number of slabs was one half of the number of interfaces. The first time that the number of slabs reduced to 1 was τ_ss_. The high cutoff density was in the range of 0.5 to 0.6 whereas the low cutoff density was in the range of 0.1 to 0.4; their precise values were selected for each system so the resulting τ_ss_ was close to the value obtained by visual inspection, which looked for a single slab in the entire simulation box.

### 2.7 Densities in the Dense and Bulk Phases

In all the simulations where a slab morphology was formed and used to calculate equilibrium properties, averages were taken over the second half of the simulations. However, when τ_ss_ occurred in the second half of a simulation, as in a few cases for LJ chains and more cases for patchy particles with large *N* and long boxes, averages were only taken from τ_ss_ to the end of the simulation, to ensure that the simulation box contained only a single slab in this calculations.

The density profile, *ρ(z),* along the *z* direction was calculated by dividing the simulation box into slices of thickness 0.1 *σ* along *z*. In each snapshot, the periodic system was translated to have the center of mass located at the center of the simulation box. The total number of particles (or beads) in each slice was then divided by its volume to yield an estimate for the density at that particular *z* in that snapshot. This estimate was then averaged over all the snapshots saved for analysis to obtain *ρ(z).* In cases where the slab was oriented with its normal along *x* or*y*(occurring only when *L_z_/L_x_* was close to 1), the density profile was calculated along that direction.

To obtain the densities, *p_d_* and *p_A_,* in the dense and bulk phases, we fit the density profile in the positive *z* range to the following function:

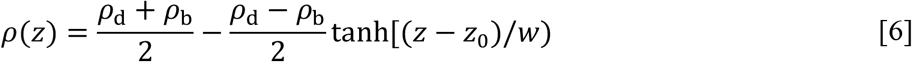

where *z_0_* represents the midpoint of the interface between the two phases, and *w* is a measure of the width of the interface.

### 2.8 Critical Temperature

The binodal, comprising bulk- and dense-phase densities as a function of temperature, was fit to the following equations:

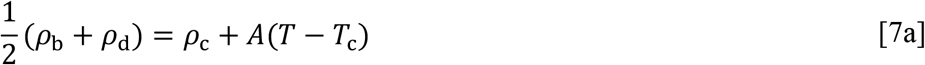

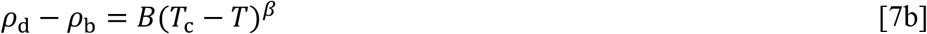

where *T_c_* is the critical temperature, *ρ_c_* is the critical density, *A* and *B* are constants, and the exponent *β* is set to 0.32.

### 2.9 Interfacial Tension

The interfacial tension *γ* was determined according to the Kirkwood-Buff method:^29^

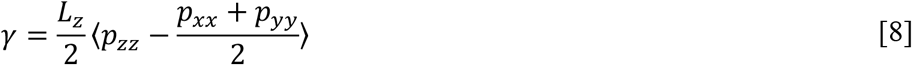

where *p_xx_*, *p_yy_*, and *p_yy_* are the diagonal elements of the pressure tensor, and the brackets indicate an equilibrium average. We calculated these diagonal elements on snapshots separated by 10 time steps for the MD simulations and averaged them over same portion of each simulation as used for calculating densities. Interfacial tension was not calculated for patchy particles, because the pressure tensor could not be properly defined due to the discontinuous nature of the interaction potential. Interfacial tension was determined for LJ particles from an MC simulation, from pressure tensor calculated at every 100 MC steps and averaged over the second half of the simulation.

### 2.10 Molecular Dynamics Simulations of Phase-Separating Tetrapeptide

We also tested SpiDec on a peptide, consisting of two copies of a phenylalanine dipeptide crosslinked at the C-termini by a disulfide bond (denoted as FFssFF), that was recently shown to phase separate.^24^ FFssFF was prepared in ChemDraw and saved in Protein Data Bank (PDB) format for force-field parametrization, which was done using Gaussian 16 at the HF/6-31G* level for atomic charges and using general Amber force field (GAFF)^30^ for other parameters. The initial configuration of 64 copies of the peptide in a water box was prepared in two steps. First, 8 copies were randomly inserted into a cubic box with a side length of 30 Å and solvated with TIP3P water^31^ using CHARMM-GUI.^32^ This system was relaxed by energy minimization (2000 steps of steepest descent and 3000 steps of conjugate gradient) and a 100 ps MD simulation at constant NVT (294 K) with a 1 fs timestep. The box with only the peptides in the last snapshot was duplicated in each of the three orthogonal directions to build a system with 64 copies in a cubic box with a side length of 60 Å, which was solvated again with 4693 TIP3P water molecules. The latter system was also relaxed by energy minimization and 500 ps of constant-NVT simulation. To further condense the system, half of the water molecules were randomly removed and the system was again relaxed by energy minimization and 500 ps of constant-NVT simulation. It was then equilibrated at constant NPT (1 bar) and with a 2 fs timestep, for 9 μs until the peptides formed a single slab with normal in the *z* direction. The side length of the cubic box at this point was reduced to 51.72 Å. Long-range electrostatic interactions were treated by the particle mesh Ewald method^33^ with nonbonded cutoff at 8 Å. Temperature was regulated by the Langevin thermostat with a damping constant of 3 ps^-1^; pressure was regulated using the Berendsen barostat.^34^ All bonds connected with hydrogen atoms were constrained using the SHAKE algorithm. ^35^

The single slab of 64 copies was finally placed in a rectangular box with *L_z_/L_x_* = 5 (*L*_x_ = 51.72 Å and *L*_z_ = 258.6 Å). To model pH 7, half of the copies were randomly chosen to have a terminal amide protonated. The system was neutralized with 32 Cl^-^ ions and solvated with 20174 TIP3P water molecules; the total number of atoms was 66986. Initial relaxation was done with energy minimization and 500 ps of constant NVT simulation with a 1 fs timestep, followed by 100 ns of equilibration at constant NPT with a 2 fs timestep. Finally constant-NVT production runs were carried out on GPUs using *pmemd.cuda*^36^ with a 2 fs timestep, at *T* = 294 K and 326 K, each for 2 μs. Snapshots were saved at 100 ps intervals for analysis.

## 3. RESULTS

### 3.1 Variety of Dense-Phase Morphologies From Spinodal Decomposition

In Fig. 2A, we display the dense-phase morphologies of LJ particles in a cubic box at *T* = 0.65, obtained from MD simulations at increasing initial densities (*ρ*_0_). The dense phase appears as a sphere at *ρ_0_* = 0.1, a cylinder at *ρ*_0_ = 0.2, a slab at *ρ*_0_ = 0.3, a hollow cylinder at *ρ*_0_ = 0.6, and a hollow sphere at *ρ*_0_ = 0.7. Similar observations are found for the other model systems, and are displayed in Fig. 2B for LJ chains at *T* = 1.7 and in Fig. S1 for HP chains at *T* = 1.05.

**Figure 2.**
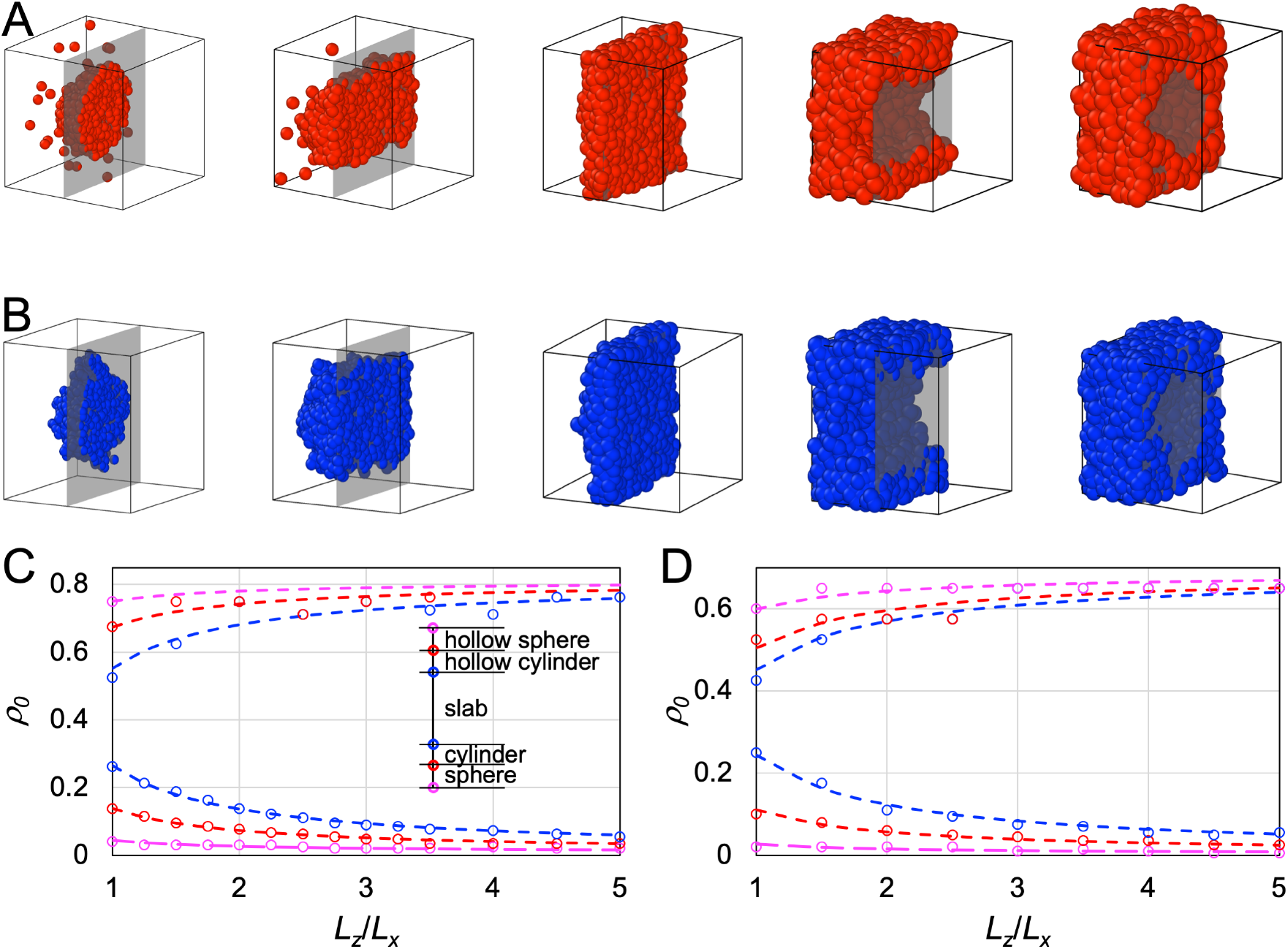
Morphologies of the dense phases of LJ particles and LJ chains over a range of initial densities inside the spinodal. (A) Morphologies for an LJ particle system of *N* = 1000 particles in a cubic box at *T* = 0.65. The dense phase appears as a sphere, cylinder, slab, hollow cylinder, and hollow sphere at *ρ*_0_ = 0.1, 0.2, 0.3, 0.6, and 0.7 respectively. The initial densities were changed by varying the box side lengths, which are shown here not to scale. At each density, the cubic box is cut by a plane (rendered as gray when the background is empty), and only the half behind the cut is displayed. (B) Corresponding results for an LJ chain system (100 10-bead chains) at *T* = 1.7 and *ρ_0_* = 0.1, 0.2, 0.3, 0.5, and 0.6. (C) Boundaries between different morphologies for LJ particles in rectangular boxes with different *L_z_*/*L_x_* ratios. The density ranges for different morphologies are illustrated in the inset. *L_x_* = 10 and *T* = 0.65. (D) Corresponding results for an LJ chain system at *L_x_* = 13 and *T* = 1.7.

We scanned the initial densities to identify the boundaries between different dense-phase morphologies. For example, for LJ particles in a cubic box (*L_z_/L_x_* = 1) at *T* = 0.65, the transitions from a single low-density phase to a spherical dense phase, from sphere to cylinder, from cylinder to slab, from slab to hollow cylinder, from hollow cylinder to hollow sphere, and from hollow sphere to a single high-density phase occur at initial densities of 0.04, 0.1375, 0.2625, 0.525, 0.675, and 0.75, respectively. Note that we checked the dense-phase morphologies on the very short timescale of phase separation via spinodal decomposition (see below for more details). On this timescale, phase separation via nucleation and growth would not have occurred. Therefore simulations from initial densities in the metastable regions, between the spinodal and binodal, would remain a single phase, and the boundaries with the two single phases effectively define the spinodal densities. In the foregoing example, the lowest and highest boundary values, 0.04 and 0.75, are the spinodal densities.

At increasing *L_z_/L_x_*, the range of initial densities for the slab morphology widens in both directions, squeezing the other inter-morphology boundaries at both ends of the range (Fig. 2C). The dependences of the boundary density values on *L_z_/L_x_* fit well to a simple function,

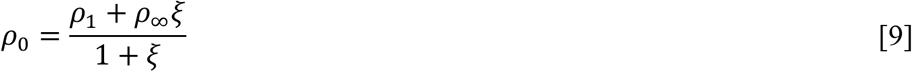

where *ξ* = L_z_/L_X_ – 1, *ρ_1_* denotes the density at *L_z_/L_X_* = 1, and *ρ_∞_* represents the density extrapolated to infinite *L_z_/L_X_*. The three lower boundaries have a common *ρ_∞_* value of 0.015, whereas the three upper boundaries have a common *ρ_∞_* value of 0.81. Similar observations are found for the other model systems. The results for LJ chains at *T* = 1.7 are shown in Fig. 2D, where the fit to eq [9] yields a *ρ_∞_* value of 0.006 for the three lower branches and a *ρ_∞_* value of 0.69 for the three upper branches. These lower and upper *ρ_∞_* values may represent the spinodal densities at infinite system sizes.

### 3.2 Slab Formation at Different *L_z_/L_x_* Ratios

The slab morphology allows easy determination of binodals (see Computational Methods). We thus pay special attention to this morphology. In Figs. 3 and 4, we display snapshots of slabs formed in simulations of the four model systems over a range of *L_z_/L_x_*. In a cubic simulation box, there is no preferred direction, and so slabs can form with the normal along any of the three orthogonal directions. The indeterminacy in slab orientation persists when *L_z_/L_x_* is slightly above 1 (Fig. 3A and 3B). Regardless of the slab orientation, the thickness of the bulk phase in simulation boxes with *L_z_/L_x_* close to 1 is small, which would make it difficult to calculate the bulk-phase density and determine the binodal.

**Figure 3.**
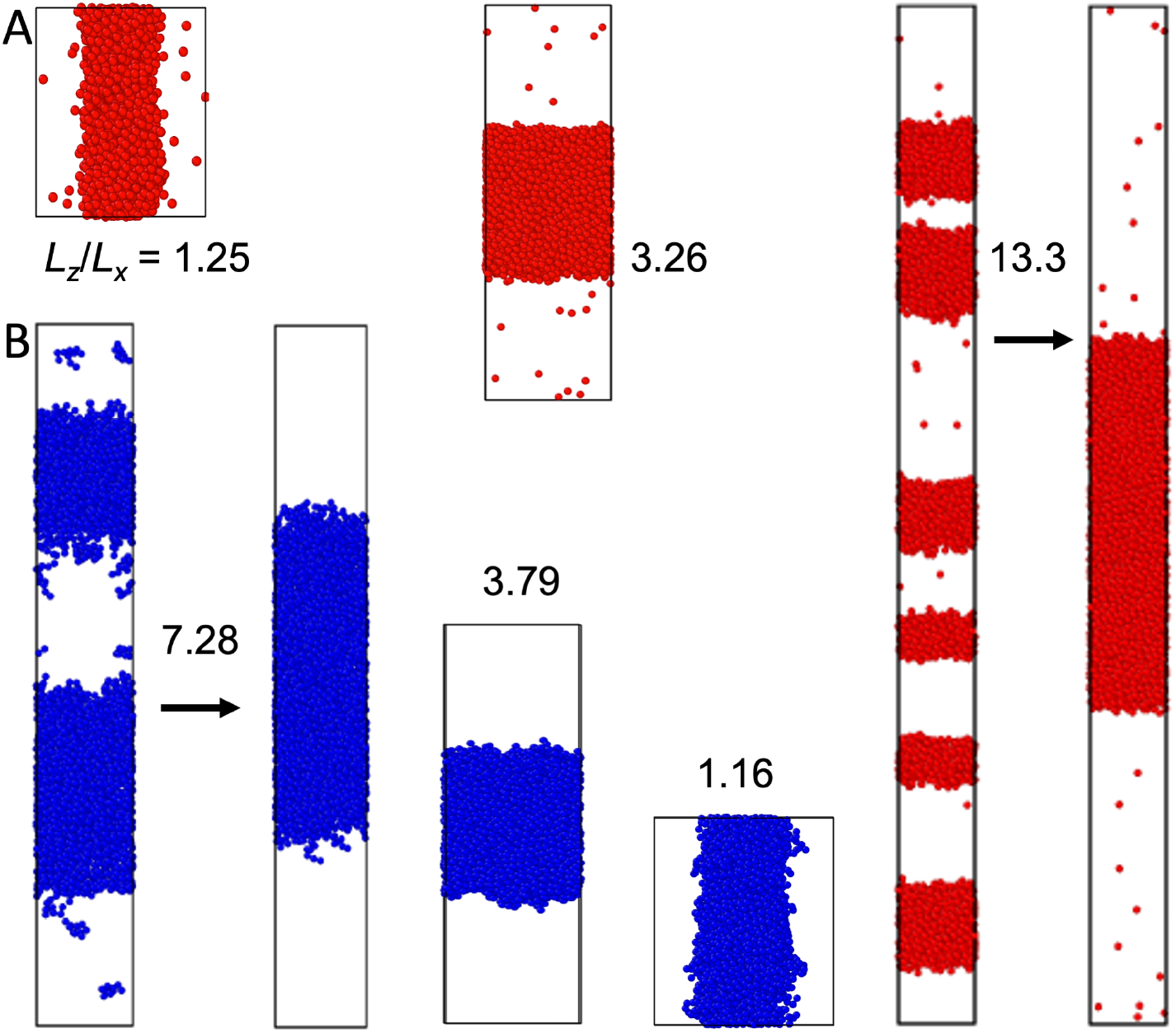
Slab formation at various *L_z_/L_x_* values. (A) LJ particle system at *N* = 4000, *T* = 0.65, *ρ*_0_ = 0.3, and *L_z_/L_x_* = 1.25, 3.26, and 13.3. (B) LJ chain system at *N* = 4000, *T* = 1.7, *ρ*_0_ = 0.25, and *L_z_/L_x_* = 1.16, 3.79, and 7.28. The *z* axis is along the vertical direction. For each system, the simulation boxes are drawn approximately to scale.

At *L_z_/L_x_* above ~2, slabs are always oriented with the normal along the *z* direction. As *L_z_/L_x_* is further increased above ~4, multiple slabs can form (Figs. 3 and 4). Over time these multiple slabs fuse one by one, eventually leading to the complete fusion into a single slab (Supporting Movie S1).

**Figure 4.**
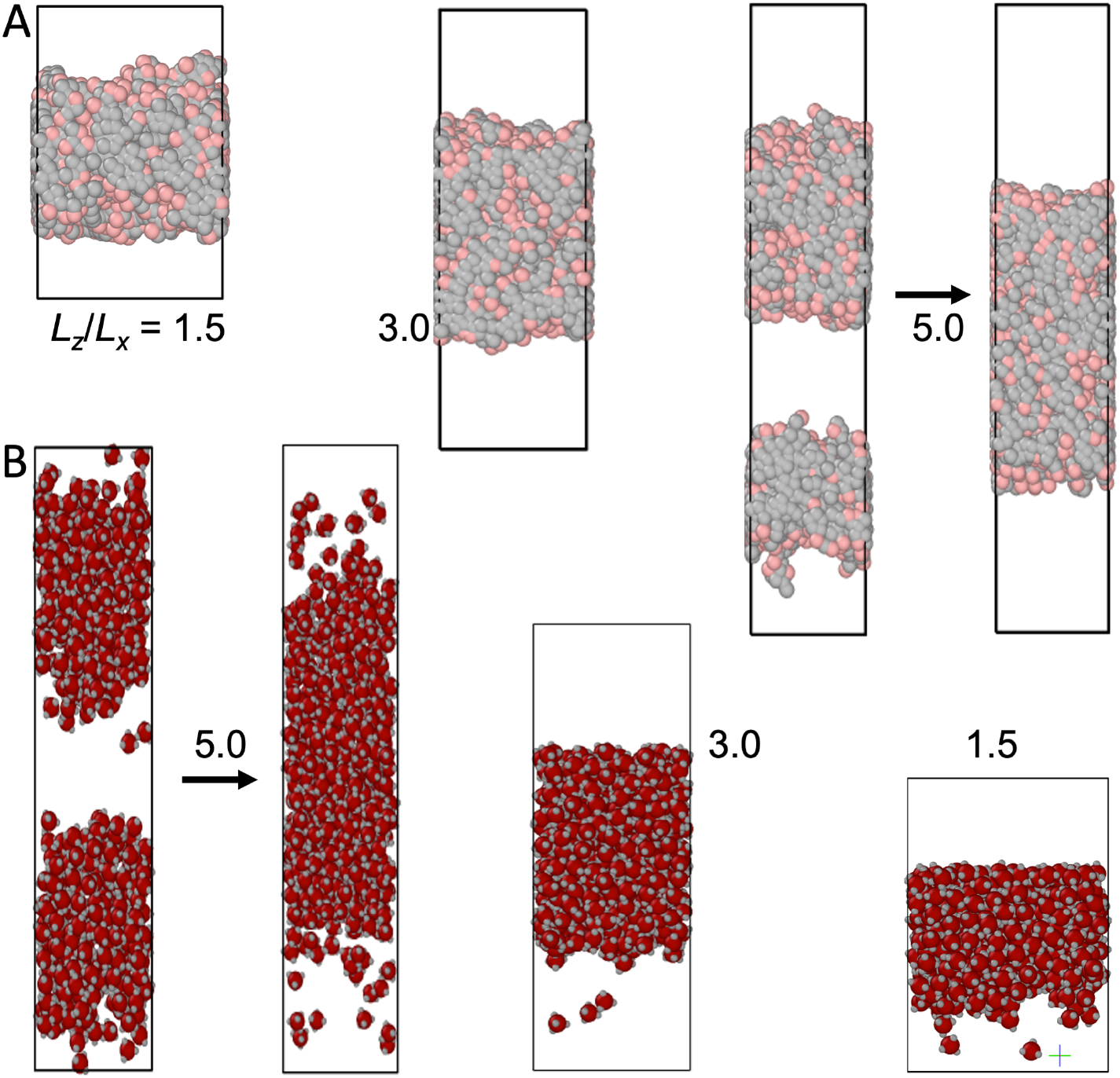
Slab formation at various *L_z_/L_x_* values. (A) HP chain system at *N* = 4000, *T* = 1.05, *ρ_0_* = 0.25, and *L_z_/L_x_* = 1.5, 3.0, and 5.0. (B) Patchy particle system at *N* = 500, *T* = 0.61, *ρ_0_* = 0.36, and *L_z_/L_x_* = 1.5, 3.0, and 5.0. The *z* axis is along the vertical direction. For each system, the simulation boxes are drawn approximately to scale.

### 3.3 Timescales for Phase Separation via Spinodal Decomposition and for Complete Slab Fusion

We devised a procedure to determine the time (τ_PS_) at which slabs first emerge from spinodal decomposition (Fig. S2A). The results for three model systems from MD simulations at *N* = 4000 are shown in Fig. S3A. For LJ particles, slab formation is quick, occurring on the order of 2 × 10^4^ time steps in MD simulations. τ_PS_ decreases with increasing *L_z_/L_x_*. For LJ and HP chains, τ_PS_ shows an even stronger initial decline (near *L_z_/L_x_* = 1). In the range of *L_z_/L_x_* from 2 to 7, the τ_PS_values (in LJ time units) for LJ and HP chains are 3-8 times as long as that for LJ particles. For patchy particles at *N*= 500 and *L_z_/L_x_* from 1.5 to 5, τ_PS_ is of the order of 3 *’* 10^5^ Monte-Carlo (MC) steps.

For the purpose of determining binodals, another important timescale is τ_ss_, for complete fusion into a single slab when multiple slabs emerge from spinodal decomposition (Fig. S2B). The results for τ_ss_ are shown in Fig. S3B. In contrast to τ_PS_, τ_ss_ exhibits a sharp increasing trend with increasing *L_z_/L_x_*. The main reason for the increase in τ_ss_ is that, with higher *L_z_/L_x_*, more slabs form initially and hence more fusion events must take place before complete fusion. A secondary reason is that, with higher *L_z_/L_x_*, slabs are farther apart initially and hence each fusion event can take longer. In any event, for LJ particles, τ_ss_ is under a million time steps even at *L_z_/L_x_* = 13.3. Since we equilibrate at least 5 million time steps anyway, so slab fusion for LJ particles at *N* = 4000 did not necessitate longer simulations. The same holds for LJ and HP chains at *N* = 4000 and *L_z_/L_x_*≤5, with τ_ss_ under 3.1 million time steps. However, for LJ chains at *L_z_/L_x_*= 7.28, τ_ss_ exceeds 5 million time steps, and as a result we extended the simulations to 200 million steps. For patchy particles at *N* = 500 and *L_z_/L_x_* from 3 to 5, τ_ss_ is 0.9 to 1.8 million MC steps.

### 3.4 Optimum in *L_z_/L_x_* and Selection of *ρ*_0_ for Binodal Determination

So it appears that there is an optimum in *L_z_/L_x_* in performing simulations to form a single slab for binodal determination. When *L_z_/L_x_* is too small (e.g., below 2), a single slab may directly form spinodal decomposition, but the orientation of the slab is indeterminant and the thickness of the bulk phase in the simulation box may be too small for a precise determination of the bulk-phase density. On the other hand, when *L_z_/L_x_* is too large (e.g., above 7), multiple slabs emerge from spinodal decomposition and can take a long time to fuse into a single slab. This long fusion time will increase the total simulation time. The optimal *L_z_/L_x_* is from 3 to 5.

Slab formation also requires a correct choice for the initial density. Fortunately, at *L_z_/L_x_* from 3 to 5, the range of initial densities leading to slab formation is pretty wide (see Fig. 2C and 2D). Here is a general procedure for selecting an appropriate *ρ*_0_ to start the simulation. The procedure involves running short simulations (e.g., 1 million steps; see Fig. S3A) at an *L_z_/L_x_* between 3 and 5 and *ρ*_0_ at *iΔρ_0_*, with *i* = 1, 2,…, and *Δρ_0_* between 0.05 and 0.1. As *ρ*_0_ is increased, a dense phase should emerge, with shapes in the order of sphere, cylinder, slab, hollow cylinder. Finally choose the midpoint in the range of initial densities leading to slab formation. This procedure can be applied once, at the lowest temperature for which the binodal is to be determined. Once the initial density is found at this temperature, it can be used for long simulations in the full temperature range to determine the binodal.

### 3.5 Binodals of Four Model Systems

Once a single slab is formed and equilibrated with the bulk phase at each temperature within a selected range, we can calculate the densities of the two phases as a function of temperature. The resulting binodals are shown in Fig. 5A and 5B for the LJ particle and LJ chain systems at *N* = 4000 and in Fig. S4A and S4B for the HP chain system at *N* = 4000 and the patchy particle system at *N* = 500. For each system, we report binodals calculated at three *L_z_/L_x_* ratios. The lowest of these ratios is close to the cubic limit, and it leads to underestimation of the critical temperature. As already pointed out, at *L_z_/L_x_* close to 1, there is very little space for the bulk phase. As the critical temperature is approached, the interface between the dense and bulk phases widens, leaving little space for a fully formed dense phase as well. As a result, the dense-phase density becomes too low whereas the bulk-phase density becomes too high. These opposite errors lead to narrowing of the binodal near the critical point and the underestimation of *T_c_*. In contrast, when *L_z_/L_x_* is within the recommended range of 3 to 5, or at a higher value as long as the single-slab morphology is sampled at equilibrium for a sufficiently long time, the calculated binodals reach convergence. Further validation of convergence is provided by a comparison of *T*_c_ values at *L_z_/L_x_* ≥ 1.66 and *N* from 1000 to 10000 for LJ particles and LJ and HP chains or *N* from 250 to 1250 for patchy particles (Fig. S5). The minimum particle number for robust binodal calculation is 4000 for LJ particles and LJ and HP chains and 500 for patchy particles. *T*_c_ values determined at smaller particle numbers show greater variations. The results for patchy particles also demonstrate that SpiDec works in MC simulations just as well as it does in MD simulations, as shown for LJ particles and LJ and HP chains. We also applied SpiDec in MC simulations of LJ particles. The binodals obtained in MD and MC simulations show close agreement (Fig. S6A).

**Figure 5.**
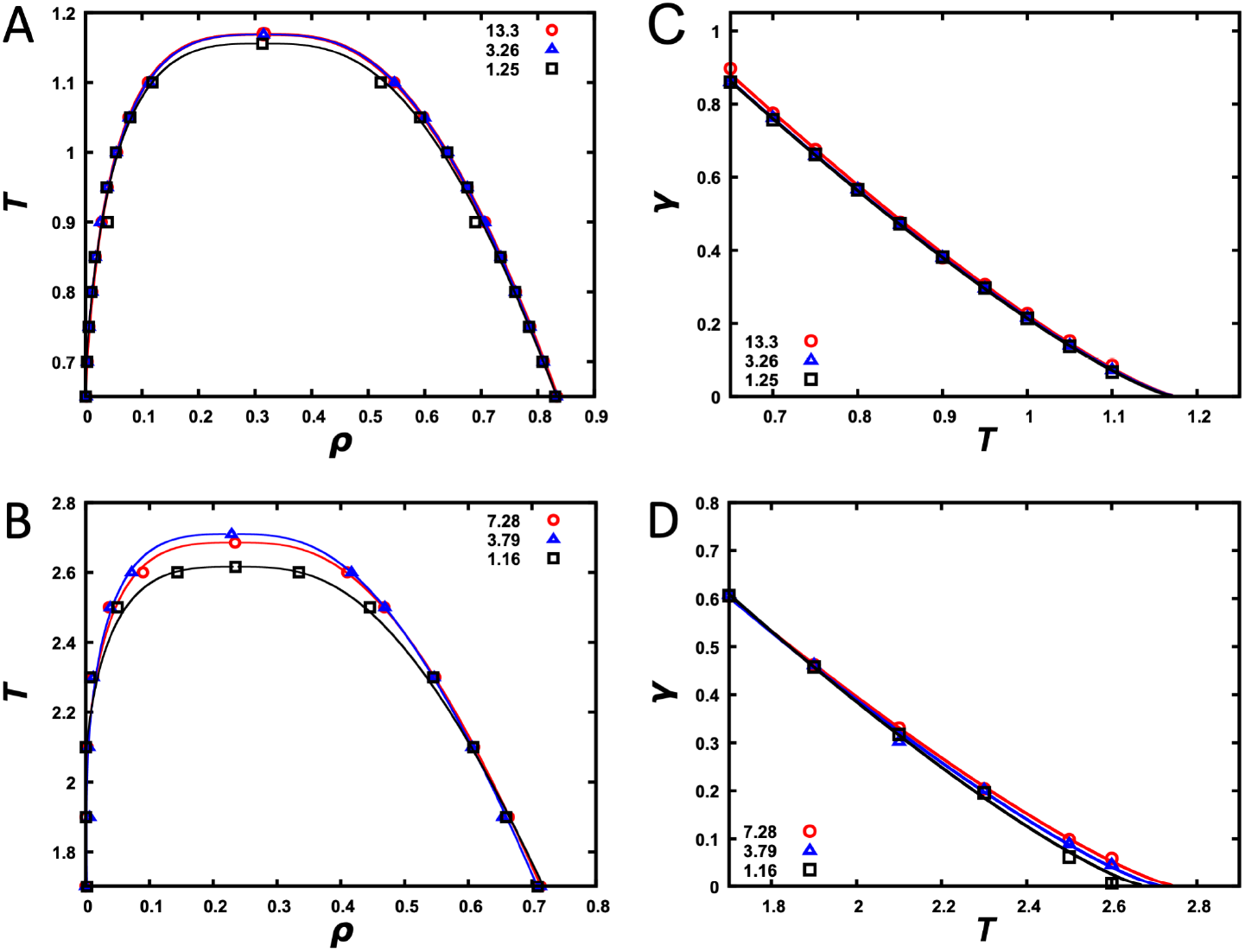
Binodals and interfacial tensions calculated from snapshots with a single slab. (A) Binodals of the LJ particle system at *N* = 4000 and *L_z_/L_x_* = 1.25, 3.26, and 13.3. (B) Binodals of the LJ chain system at *N* = 4000 and *L_z_/L_x_* = 1.16, 3.79, and 7.28. (C) Interfacial tension versus temperature for the LJ particle system at the three *L_z_*/*L_x_* ratios. (D) Interfacial tension versus temperature for LJ chain system at the three *L*_z_/*L_x_* ratios.

### 3.6 Interfacial Tensions of Three Model Systems

The single-slab morphology generated by SpiDec also allows us to calculate the interfacial tension. The results are shown in Fig. 5C and 5D for the LJ particle and LJ chain systems and in Fig. S4C for the HP chain system, all at *N* = 4000. Again, as long as *L_z_*/*L_x_* is above 2, convergent results are obtained. We also obtained very similar interfacial tension for the LJ particle system from both MD and MC simulations (Fig. S6B).

### 3.6 Application of SpiDec to a System Modeled at the All-Atom Level in Explicit Solvent

Finally we tested SpiDec on the tetrapeptide FFssFF (Fig. 6A) that was recently shown to phase separate.^24^ Starting from a random, loose configuration solvated in water, 64 copies of the peptide condensed into a single slab (Fig. 6B). We subsequently solvated this single slab in an elongated box (*L_z_*/*L_x_* = 5). The copies equilibrated between the dense and bulk phases. We calculated the average concentrations (expressed as wt/wt, i.e., weight of peptide over weight of water) along the *z* axis, and fit the concentration profile to eq [6] to obtain the dense- and bulk-phase concentrations (Fig. 6C). At 294 K, the two concentrations are 0.94 wt/wt and 0.0085 wt/wt, respectively. These values are each about 3-fold higher than the experimental counterparts,^24^ reflecting the need for improved force-field parameterization. We also carried out a simulation at 326 K. At the elevated temperature, the two densities move toward each other (Fig. 6D), as expected for a system showing upper critical solution temperature.

**Figure 6.**
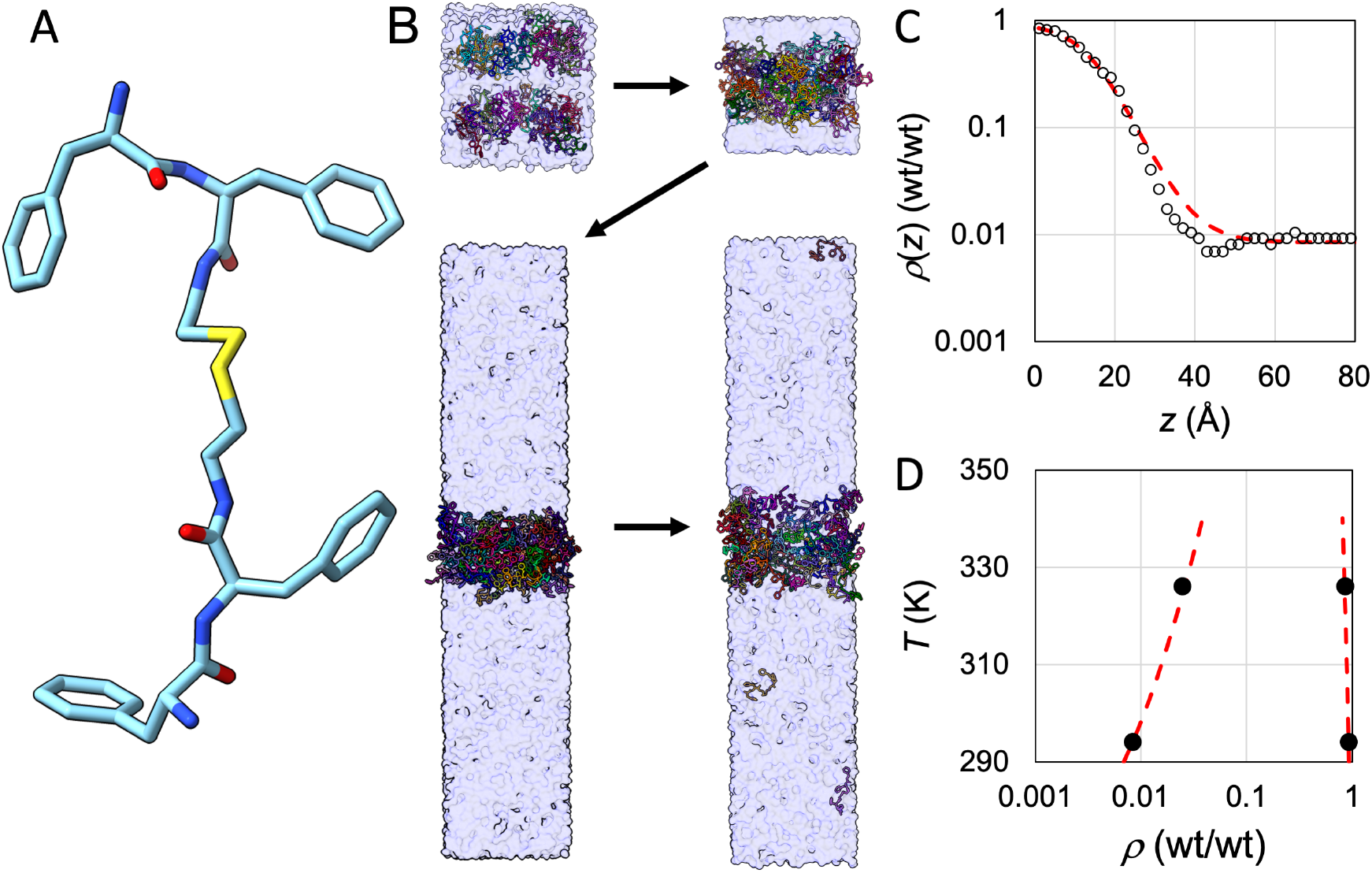
Application of SpiDec to a phase-separating peptide. (A) Structure of FFssFF. (B) The SpiDec simulation procedure. First, a random, loose configuration condensed into a single slab. Then the slab was solvated into an elongated box and the two phases reached equilibrium. The fourth snapshot shown was taken at 1.065 μs of a 2-μs simulation at 294 K. (C) Concentration profile (circles) at 294 K and fit to eq [6] (red curve). Concentrations were calculated by counting copy numbers in 2-Å slices along the *z* direction; the location of each copy was represented by the midpoint of the central disulfide bond. The conversion of concentrations to wt/wt used a molecular weight of 741.5 Da for the peptide and a density of 1 kg/L for water. (D) Dilute- and dense-phase concentrations at two temperatures, 294 K and 326 K. The red curve shows a binodal to guide the eye.

## 4. DISCUSSION

We have characterized the morphologies and timescales of spinodal decomposition in model systems and demonstrated that slab formation via spinodal decomposition can be used to calculate the binodal and interfacial tension. At different initial densities in a rectangular simulation box, spinodal decomposition produces distinct dense-phase morphologies, including sphere, cylinder, slab, hollow cylinder, and hollow sphere. The range of initial densities for slab formation is the widest among all the dense-phase morphologies, and this range further widens as *L_z_/L_x_* increases. At *L_z_/L_x_* above ~3, multiple slabs emerge from spinodal decomposition. The time for multiple slabs to fuse into a single slab increases rapidly with increasing *L_z_/L_x_*. The optimal *L_z_/L_x_* is thus 3 to 5 for computing the binodal and interfacial tension. Most importantly, we have shown that the SpiDec method is effective both for model systems and for all-atom phase-separating peptides solvated in TIP3P water.

In addition to computing the equilibrium properties (binodal and interfacial tension) of phase separation, SpiDec simulations can be adapted to study dynamic properties of related processes. In particular, the fusion between multiple slabs observed here corresponds to condensate fusion. Therefore SpiDec simulations at large *L_z_/L_x_* can be used to model condensate coarsening, potentially providing a molecular view into the mechanism and kinetics of the coarsening process. Moreover, all the simulations reported above have resulted in complete phase separation, but at lower temperatures (or equivalently, at stronger intermolecular attraction), as expected,^37^ spinodal decomposition may be arrested, leading to gelation (Supporting Movie S2). Therefore SpiDec simulations at those conditions provide a means to study condensate gelation.

For the phase-separating tetrapeptide, our simulation results agree qualitatively with experimental data,^24^ but quantitatively, the computed concentrations in the two phases are each about 3-fold higher than the experimental counterparts. Binodals are exquisitely sensitive to force fields, and now, with SpiDec, we will have the opportunity to use experimental binodals for force-field parameterization of IDPs, leading to accurate modeling of IDPs and their condensate properties.

## ASSOCIATED CONTENT

### Supporting Information

The following Supporting Information is available free of charge:

Computational methods section; six additional figures (Figures S1 to S6) presenting the dense-phase morphologies of HP chains; the timescales for phase separation and for complete fusion of multiple slabs into a single slab, and illustration of their determination; binodal and interfacial tension of HP chains and binodal of patchy particles; the effects of system size and *L_z_/L_x_* ratio on the calculated critical temperature; comparison of the binodal and interfacial tension of LJ particles determined by MD and MC simulations; and Movie S1 showing the phase separation of LJ particles via spinodal decomposition and the fusion of multiple slabs, and Movie S2 showing arrested spinodal decomposition leading to gelation of HP chains at *T* = 0.2.

## AUTHOR INFORMATION

### Author Contributions

KM and HXZ designed research; KM and RP carried out research and analyzed data. KM and HXZ wrote the manuscript.

### Notes

The authors declare no competing financial interest.

## ACKNOWLEDGMENT

This work was supported by National Institutes of Health Grant GM118091.

## ABBREVIATIONS

HP: hydrophobic-hydrophilic
IDP: intrinsically disordered protein
LJ: Lennard-Jones
MC: Monte Carlo
MD: molecular dynamics
WCA: Weeks-Chandler-Anderson

## REFERENCES

1. Thomson, J. A.; Schurtenberger, P.; Thurston, G. M.; Benedek, G. B. Binary liquid phase separation and critical phenomena in a protein/water solution. Proc Natl Acad Sci U S A 1987, 84, 7079–7083.

2. Bracha, D.; Walls, M. T.; Wei, M.-T.; Zhu, L.; Kurian, M.; Avalos, J. L.; Toettcher, J. E.; Brangwynne, C. P. Mapping Local and Global Liquid Phase Behavior in Living Cells Using Photo-Oligomerizable Seeds. Cell 2018, 175, 1467–1480.e13.

3. Kasinsky, H. E.; Gowen, B. E.; Ausió, J. Spermiogenic Chromatin Condensation Patterning in Several Hexapods May Involve Phase Separation Dynamics by Spinodal Decomposition or Microemulsion Inversion (Nucleation). Tissue Cell 2021, 73, 101648.

4. Zeng, X.; Holehouse, A. S.; Chilkoti, A.; Mittag, T.; Pappu, R. V. Connecting Coil-to-Globule Transitions to Full Phase Diagrams for Intrinsically Disordered Proteins. Biophys J 2020, 119, 402–418.

5. Binder, K.; Block, B. J.; Virnau, P.; Tröster, A. Beyond the Van Der Waals Loop: What Can Be Learned from Simulating Lennard-Jones Fluids Inside the Region of Phase Coexistence. Am J Phys 2012, 80, 1099–1109.

6. Díaz-Herrera, E.; Cerón-García, E.; Bryan Gutiérrez, A.; Chapela, G. A. Finite Size Effect on the Existence of the Liquid–Vapour Spinodal Curve. Mol Phys 2022, 120, e1989071.

7. Mazarakos, K.; Qin, S.; Zhou, H. X. Calculating Binodals and Interfacial Tension of Phase-Separated Condensates from Molecular Simulations, with Finite-Size Corrections. Methods Mol Biol 2022, in press.

8. Rao, M.; Levesque, D. Surface Structure of a Liquid Film. J Chem Phys 1976, 65, 3233–3236.

9. Mazarakos, K.; Zhou, H. X. Macromolecular Regulators Have Matching Effects on the Phase Equilibrium and Interfacial Tension of Biomolecular Condensates. Protein Sci 2021, 30, 1360–1370.

10. Mazarakos, K.; Zhou, H. X. Multiphase organization is a second phase transition within multi-component biomolecular condensates. J Chem Phys 2022, 156, 191104.

11. Das, S.; Amin, A. N.; Lin, Y.-H.; Chan, H. S. Coarse-Grained Residue-Based Models of Disordered Protein Condensates: Utility and Limitations of Simple Charge Pattern Parameters. Phys Chem Chem Phys 2018, 20, 28558–28574.

12. Dignon, G. L.; Zheng, W.; Kim, Y. C.; Best, R. B.; Mittal, J. Sequence Determinants of Protein Phase Behavior From a Coarse-Grained Model. PLoS Comput Biol 2018, 14, e1005941.

13. Statt, A.; Casademunt, H.; Brangwynne, C. P.; Panagiotopoulos, A. Z. Model for Disordered Proteins with Strongly Sequence-Dependent Liquid Phase Behavior. J Chem Phys 2020, 152, 075101.

14. Tang, Y.; Bera, S.; Yao, Y.; Zeng, J.; Lao, Z.; Dong, X.; Gazit, E.; Wei, G. Prediction and Characterization of Liquid-Liquid Phase Separation of Minimalistic Peptides. Cell Rep Phys Sci 2021, 2, 100579.

15. Zheng, W.; Dignon, G. L.; Jovic, N.; Xu, X.; Regy, R. M.; Fawzi, N. L.; Kim, Y. C.; Best, R. B.; Mittal, J. Molecular Details of Protein Condensates Probed by Microsecond Long Atomistic Simulations. J Phys Chem B 2020, 124, 11671–11679.

16. Welsh, T. J.; Krainer, G.; Espinosa, J. R.; Joseph, J. A.; Sridhar, A.; Jahnel, M.; Arter, W. E.; Saar, K. L.; Alberti, S.; Collepardo-Guevara, R., et al. Surface Electrostatics Govern the Emulsion Stability of Biomolecular Condensates. Nano Letters 2022, 22, 612–621.

17. Panagiotopoulos, A. Z. Direct Determination of Phase Coexistence Properties of Fluids by Monte Carlo Simulations in a New Ensemble. Mol Phys 1987, 61, 813–826.

18. Ghosh, A.; Mazarakos, K.; Zhou, H. X. Three Archetypical Classes of Macromolecular Regulators of Protein Liquid-Liquid Phase Separation. Proc Natl Acad Sci U S A 2019, 116, 19474–19483.

19. Nguemaha, V.; Zhou, H. X. Liquid-Liquid Phase Separation of Patchy Particles Illuminates Diverse Effects of Regulatory Components on Protein Droplet Formation. Sci Rep 2018, 8, 6728.

20. Li, L.; Srivastava, S.; Andreev, M.; Marciel, A. B.; de Pablo, J. J.; Tirrell, M. V. Phase Behavior and Salt Partitioning in Polyelectrolyte Complex Coacervates. Macromolecules 2018, 51, 2988–2995.

21. Lytle, T. K.; Radhakrishna, M.; Sing, C. E. High Charge Density Coacervate Assembly via Hybrid Monte Carlo Single Chain in Mean Field Theory. Macromolecules 2016, 49, 9693–9705.

22. McCarty, J.; Delaney, K. T.; Danielsen, S. P. O.; Fredrickson, G. H.; Shea, J. E. Complete Phase Diagram for Liquid-Liquid Phase Separation of Intrinsically Disordered Proteins. J Phys Chem Lett 2019, 10, 1644–1652.

23. Qin, S.; Zhou, H. X. Fast Method for Computing Chemical Potentials and Liquid-Liquid Phase Equilibria of Macromolecular Solutions. J Phys Chem B 2016, 120, 8164–74.

24. Abbas, M.; Lipiński, W. P.; Nakashima, K. K.; Huck, W. T. S.; Spruijt, E. A Short Peptide Synthon for Liquid–Liquid Phase Separation. Nat Chem 2021, 13, 1046–1054.

25. Mazarakos, K.; Zhou, H.-X. Multiphase Organization Is a Second Phase Transition Within Multi-Component Biomolecular Condensates. J Chem Phys 156, DOI: 10.1063/5.0088004.

26. Weeks, J. D.; Chandler, D.; Andersen, H. C. Role of Repulsive Forces in Determining the Equilibrium Structure of Simple Liquids. J Chem Phys 1971, 54, 5237–5247.

27. Kern, N.; Frenkel, D. Fluid–Fluid Coexistence in Colloidal Systems with Short-Ranged Strongly Directional Attraction. J Chem Phys 2003, 118, 9882–9889.

28. Glaser, J.; Nguyen, T. D.; Anderson, J. A.; Lui, P.; Spiga, F.; Millan, J. A.; Morse, D. C.; Glotzer, S. C. Strong Scaling of General-Purpose Molecular Dynamics Simulations on GPUs. Comput Phys Commun 2015, 192, 97–107.

29. Kirkwood, J. G.; Buff, F. P. The Statistical Mechanical Theory of Surface Tension. J Chem Phys 1949, 17, 338–343.

30. Wang, J.; Wolf, R. M.; Caldwell, J. W.; Kollman, P. A.; Case, D. A. Development and Testing of a General Amber Force Field. J Comput Chem 2004, 25, 1157–1174.

31. Jorgensen, W. L.; Chandrasekhar, J.; Madura, J. D.; Impey, R. W.; Klein, M. L. Comparison of Simple Potential Functions for Simulating Liquid Water. J Chem Phys 1983, 79, 926–935.

32. Jo, S.; Kim, T.; Iyer, V. G.; Im, W. CHARMM-GUI: A Web-Based Graphical User Interface for CHARMM. J Comput Chem 2008, 29, 1859–1865.

33. Essmann, U.; Perera, L.; Berkowitz, M. L.; Darden, T.; Lee, H.; Pedersen, L. G. A Smooth Particle Mesh Ewald Method. J Chem Phys 1995, 103, 8577–8593.

34. Berendsen, H. J. C.; Postma, J. P. M.; Gunsteren, W. F. v.; DiNola, A.; Haak, J. R. Molecular Dynamics with Coupling to an External Bath. J Chem Phys 1984, 81, 3684–3690.

35. Ryckaert, J.-P.; Ciccotti, G.; Berendsen, H. J. C. Numerical Integration of the Cartesian Equations of Motion of a System with Constraints: Molecular Dynamics of n-Alkanes. J Comput Phys 1977, 23, 327–341.

36. Salomon-Ferrer, R.; Götz, A. W.; Poole, D.; Le Grand, S.; Walker, R. C. Routine Microsecond Molecular Dynamics Simulations with AMBER on GPUs. 2. Explicit Solvent Particle Mesh Ewald. J Chem Theory Comput 2013, 9, 3878–3888.

37. Foffi, G.; Michele, C. D.; Sciortino, F.; Tartaglia, P. Arrested Phase Separation in a Short-Ranged Attractive Colloidal System: A Numerical Study. J Chem Phys 2005, 122, 224903.

