## Supporting Information for "SpiDec: Computing Binodals and Interfacial Tension of Biomolecular Condensates From Simulations of Spinodal Decomposition"

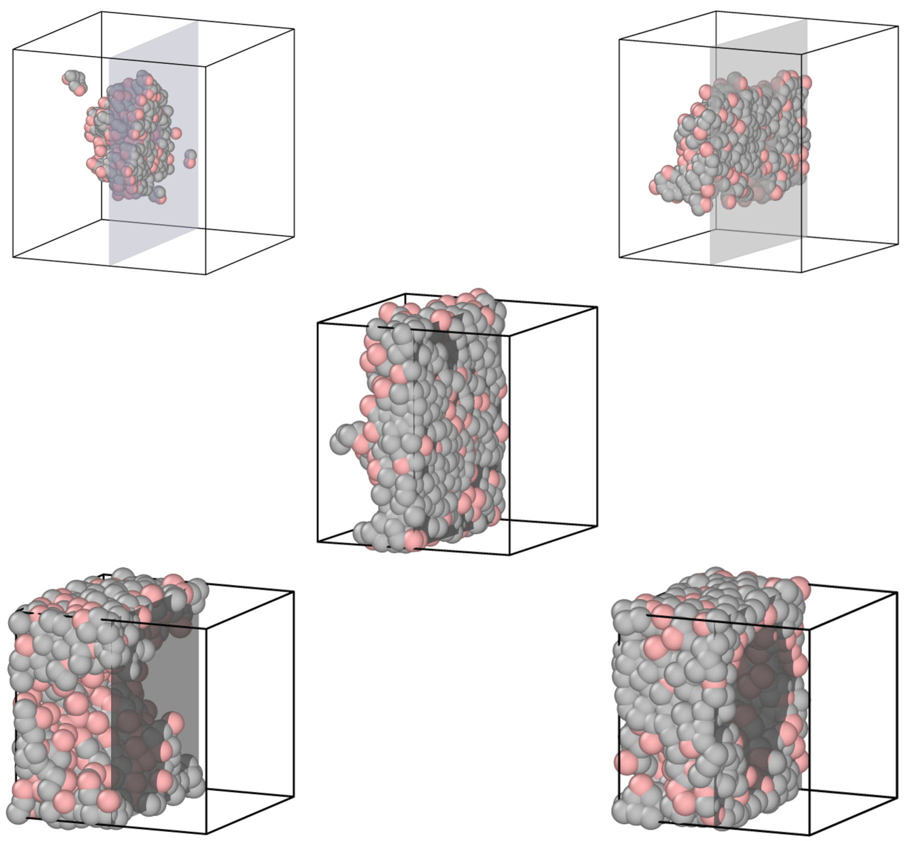

**Figure S1.** Morphologies of the dense phases of HP chains over a range of initial densities inside the spinodal. 100 chains of 10 beads each were simulated in a cubic box at $T = 1.05$. The dense phase appears as a sphere, cylinder, slab, hollow cylinder, and hollow sphere at $\rho_0 =$ 0.05, 0.1, 0.3, 0.4, and 0.5 respectively. The initial densities were changed by varying the box side lengths, which are shown here not to scale. At each density, the cubic box is cut by a plane (rendered as gray when the background is empty), and only the half behind the cut is displayed.

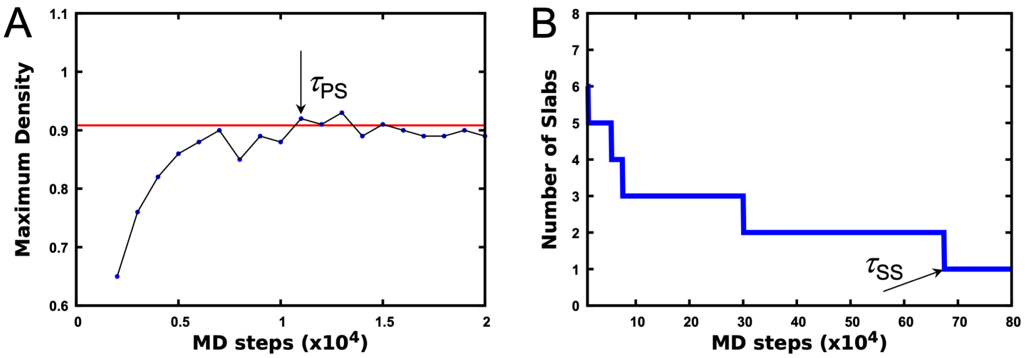


**Figure S2.** Determination of the timescales for phase separation and for complete fusion of multiple slabs into a single slab, illustrated by the LJ particle system at $N = 4000$, $T = 0.65$, $\rho_0 = 0.3$, and $L_z/L_x = 13.3$. (A) Maximum density within a snapshot, as a function of time. A horizontal line indicates the long-time plateau of the maximum density. An arrow indicates the first snapshot where the maximum density exceeds the long-time plateau, defining the timescale for phase separation. Here the thickness of the slices for density calculation was $1\sigma$, which was used in all the model systems except for HP chains. In that case the thickness was increased to $10\sigma$, because spinodal decomposition produced a domain with a thick, dense core but without flat interfaces; that core would be counted as a slab if the slice thickness were $1\sigma$. (B) Number of slabs as a function of time. An arrow indicates the first snapshot where the number of slabs reaches 1, defining the timescale for complete fusion into a single slab.

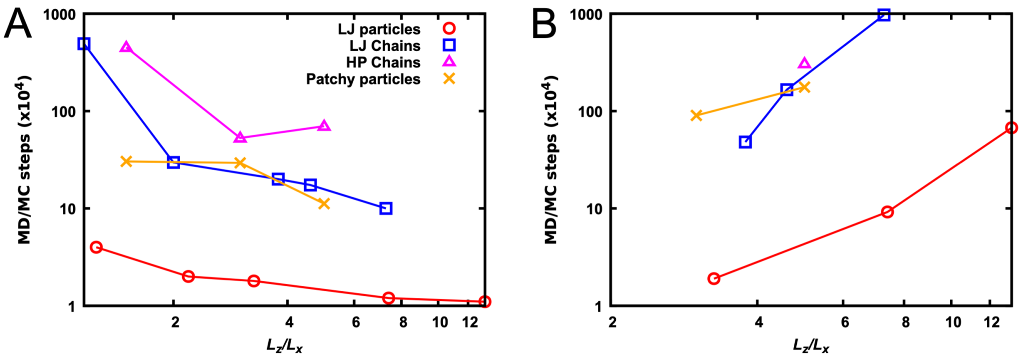


**Figure S3.** Times for phase separation and for complete fusion into a single slab. (A) Time for slabs for first emerge from spinodal decomposition. (B) Time for the complete fusion of multiple slabs into a single slab. For LJ particles and LJ and HP chains, the particle (bead) numbers are all 4000; the temperatures are 0.65, 1.7, 1.05, respectively; and the initial densities are 0.3, 0.25, 0.25, respectively. The parameters for patchy particles are $N = 500$, $T = 0.61$, and $\rho_0 = 0.36$. Times are shown in units of $10^4$ MD or MC steps.

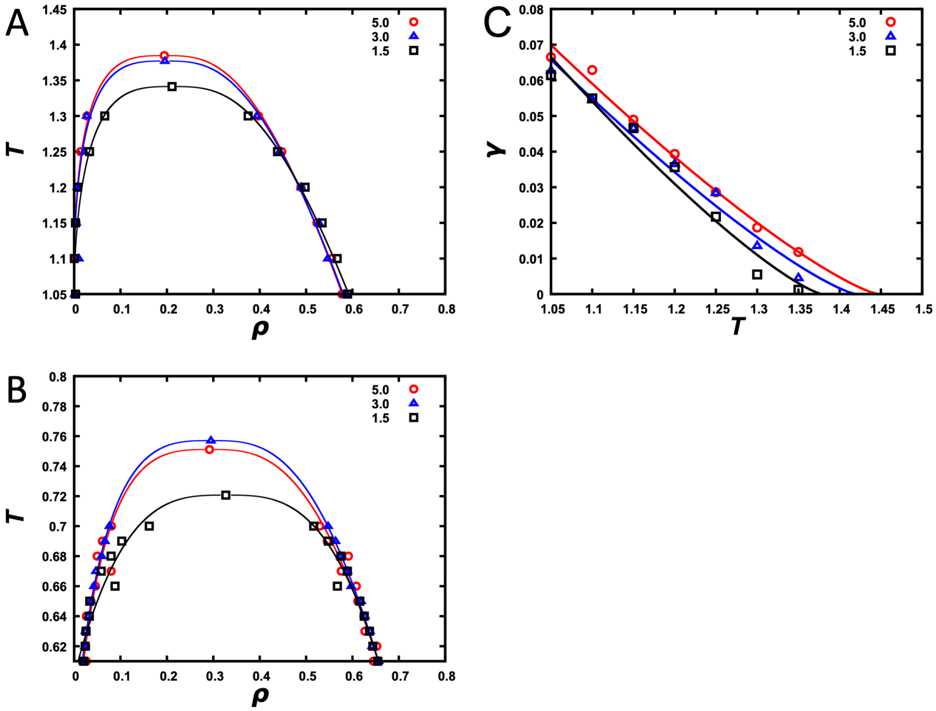


**Figure S4.** Binodals and interfacial tensions calculated from snapshots with a single slab. (A) Binodals of the HP chain system at $N = 4000$ and $L_z/L_x = 1.5$, 3.0, and 5.0. (B) Binodals of the patchy particle system at $N = 500$ and $L_z/L_x = 1.5$, 3.0, and 5.0. (C) Interfacial tension versus temperature for the HP chain system at the three $L_z/L_x$ ratios.

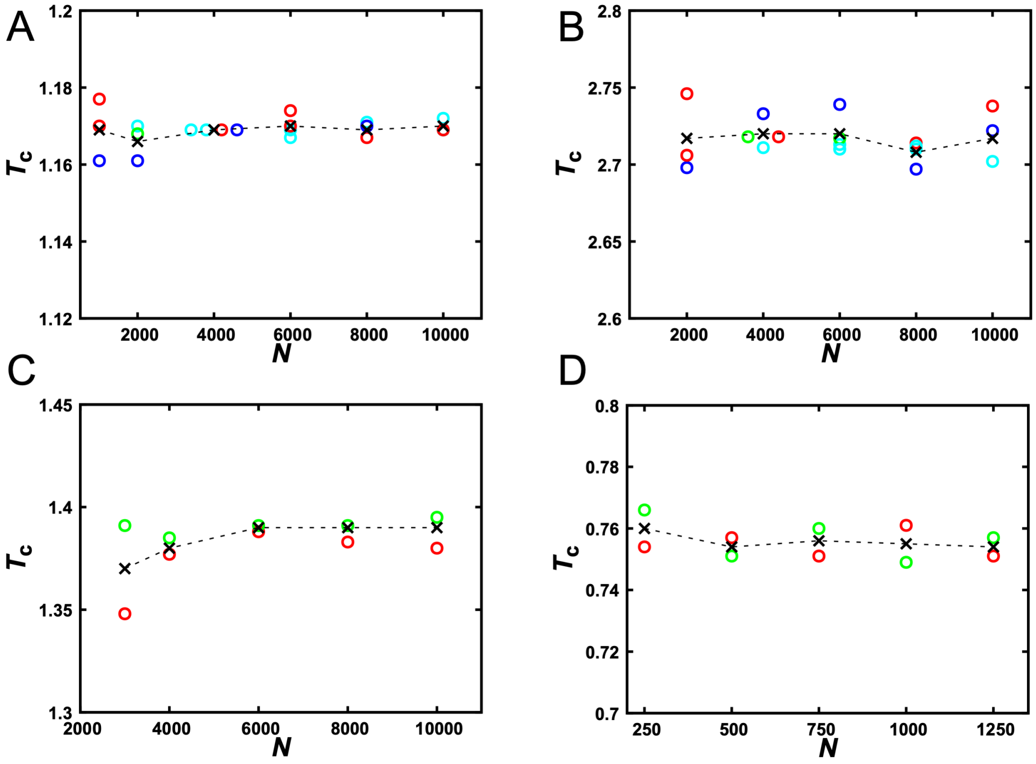


**Figure S5.** Comparison of critical temperatures obtained at various $N$ values and $L_z/L_x$ ratios. (A) LJ particles. (B) LJ chains. (C) HP chains. (D) Patchy particles. $T_c$ data are displayed in circles with the following color scheme according to $L_z/L_x$ ratios: blue, 1.66-2.5; red, 2.5-4.0; green, 4.0-6.0; and cyan, > 6.0. Crosses show averages $T_c$ values at a given $N$.

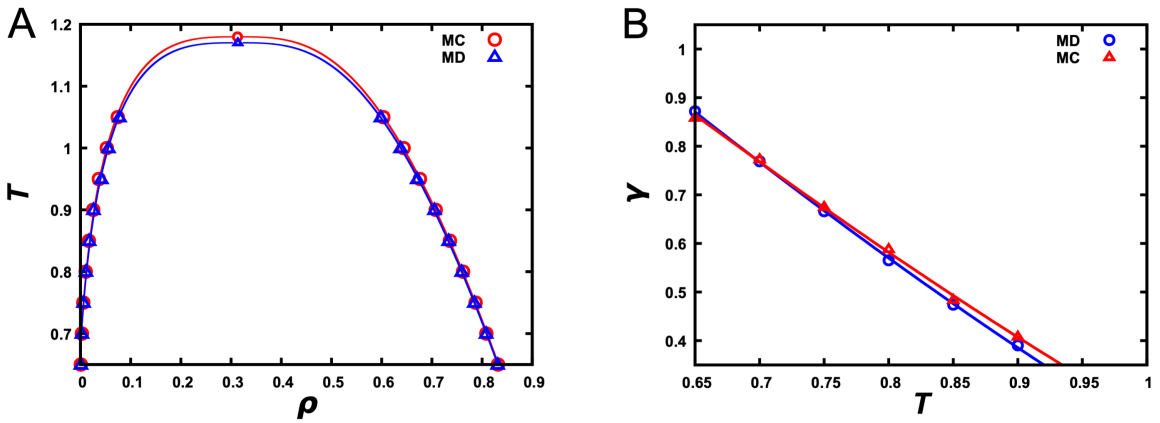


**Figure S6.** Comparison of binodals and interfacial tensions calculated from MD and MC simulations of LJ particles at $N = 1000$ and $L_z/L_x = 3$. (A) Binodals. (B) Interfacial tensions.

**Supporting Movie S1.** Phase separation of LJ particles via spinodal decomposition and the fusion of multiple slabs, at $N = 4000$, $T = 0.65$, $\rho_0 = 0.3$, and $L_z/L_x = 13.3$. The entire movie consists of 800 snapshots sampled at intervals of 1000 MD steps. Snapshots 9 and 800 are shown in Fig. 2A.

**Supporting Movie S2.** Arrested spinodal decomposition and gelation of HP chains, at $N = 4000$, $T = 0.2$, $\rho_0 = 0.25$, and $L_z/L_x = 5$. The entire movie consists of 400 snapshots sampled at intervals of 10000 MD steps.
